# Mechanical dysfunction induced by a hypertrophic cardiomyopathy mutation is the primary driver of cellular adaptation

**DOI:** 10.1101/2020.05.04.067181

**Authors:** Sarah R. Clippinger, Paige E. Cloonan, Wei Wang, Lina Greenberg, W. Tom Stump, Paweorn Angsutararux, Jeanne M. Nerbonne, Michael J. Greenberg

## Abstract

Familial hypertrophic cardiomyopathy (HCM), a leading cause of sudden cardiac death, is primarily caused by mutations in sarcomeric proteins. The pathogenesis of HCM is complex, with functional changes that span scales from molecules to tissues. This makes it challenging to deconvolve the biophysical molecular defect that drives the disease pathogenesis from downstream changes in cellular function. Here, we examined a HCM mutation in troponin T, R92Q. We demonstrate that the primary molecular insult driving the disease pathogenesis is mutation-induced alterations in tropomyosin positioning, which causes increased molecular and cellular force generation during calcium-based activation. We demonstrate computationally that these increases in force are direct consequences of the initial molecular insult. This altered cellular contractility causes downstream alterations in gene expression, calcium handling, and electrophysiology. Taken together, our results demonstrate that molecularly driven changes in mechanical tension drive the early disease pathogenesis, leading to activation of adaptive mechanobiological signaling pathways.

## Introduction

Hypertrophic cardiomyopathy (HCM) is the leading cause of sudden cardiac death in people under age 30. HCM is characterized by hypertrophy of the left ventricular wall and the intraventricular septum, myocyte disarray, fibrosis, and diastolic dysfunction. HCM is also associated with marked alterations in cardiomyocyte functioning, including changes in electrophysiology, contractility and calcium handling (1). Large scale sequencing of families has revealed that HCM is caused by mutations in sarcomeric proteins involved in cardiac contraction, including troponin T (2).

Disease presentation in HCM is quite complex, with functional differences seen at scales ranging from molecules to tissues; however, at a fundamental level, the molecular trigger that drives the disease pathogenesis is alterations in the abundance, stability, and/or functioning of the mutant protein (3). This initial trigger leads to downstream adaptive and maladaptive processes, some of which can take years to decades to manifest, including ventricular remodeling, and eventually symptomatic cardiac dysfunction. Given the inherent complexity of HCM, it has been challenging to link the molecular and cellular phenotypes and to dissect the initial biophysical trigger from secondary adaptive processes.

To better understand the connection between the initial molecular insult and cellular dysfunction in the early disease pathogenesis of HCM, we examined a point mutation in troponin T, R92Q (Fig. 1A), identified in several unrelated families, that causes pronounced ventricular hypertrophy and a relatively high incidence of sudden cardiac death (2). R92Q has been studied in several model systems, including feline (4) and rat (5) cardiomyocytes, rabbit skeletal myofibrils (6), quail myotubes (7), and transgenic mice (8). These studies have resulted in conflicting conclusions about the effects of the mutation, at least in part due to phenotypic differences between species. For example, the widely studied transgenic mouse model of R92Q (8) recapitulates some, but not all, aspects of the disease phenotypes seen in humans. Elegant experiments by the Tardiff lab have shown that the disease presentation in mice depends on the myosin heavy chain isoform expressed, with different phenotypes seen when using the faster (*MYH6*) isoform found in mouse ventricles or the slower (*MYH7*) isoform found in human ventricles (9). These studies highlight the need to study the mutation in humanized systems.

**Figure 1.**
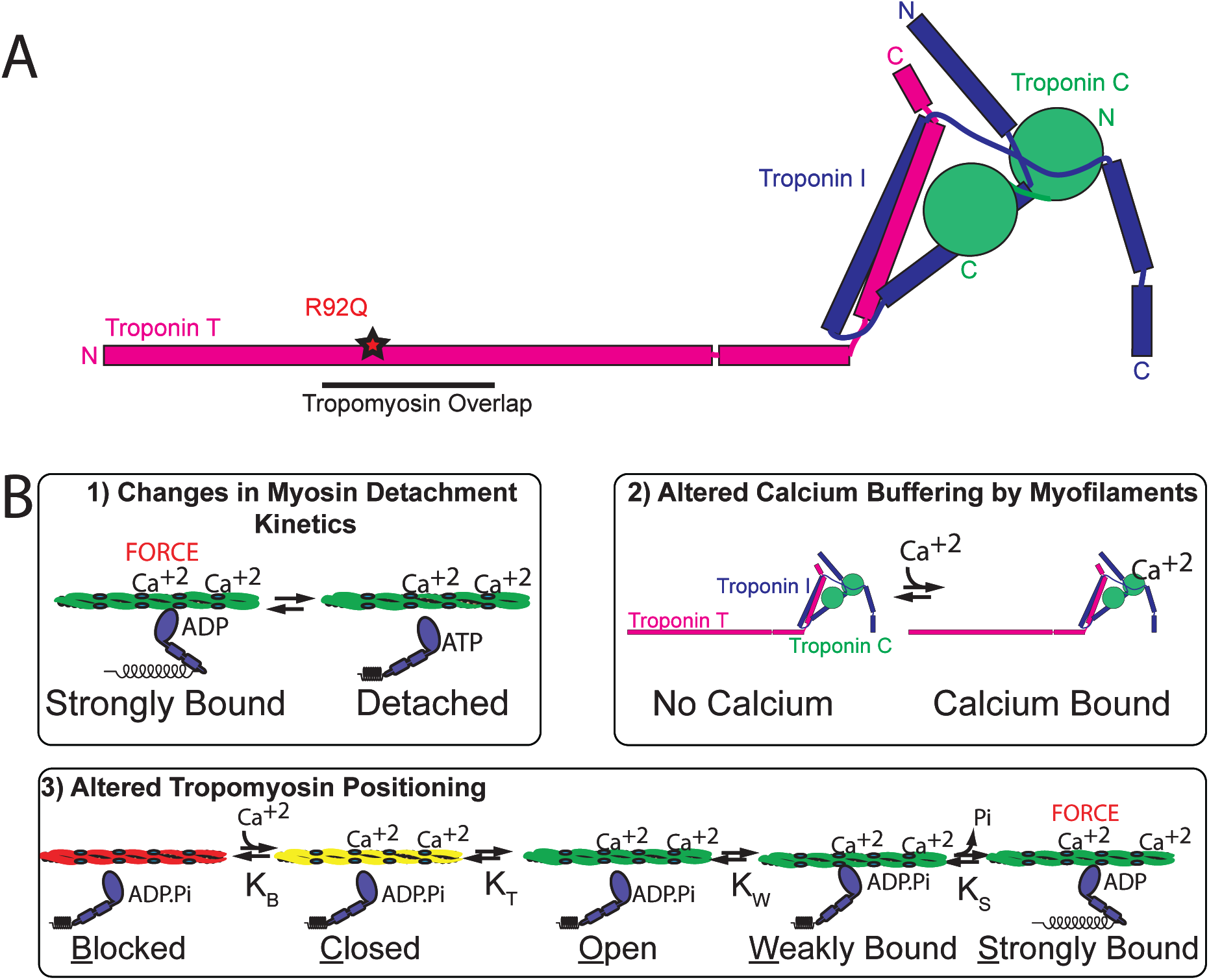
R92Q mutation in troponin T causes hypertrophic cardiomyopathy. (A) Cartoon of the troponin complex based on (70). R92Q is located in the region of troponin T that interacts with tropomyosin, near the tropomyosin overlap region. (B) Models for the molecular mechanism of R92Q.

Troponin T is part of the troponin complex, which, together with tropomyosin, regulates the calcium-dependent interactions between myosin and the thin filament that drive muscle contraction. Three models have been put forward to describe the initial molecular insult that drives the disease pathogenesis of R92Q (Fig. 1B). 1) R92Q could affect the cycling kinetics of myosins that are bound to the thin filament (9). In this model, one would expect to observe a change in the amount of time that myosin remains bound to the thin filament during crossbridge cycling in the mutant. 2) R92Q could increase the calcium affinity of the troponin complex, leading to altered calcium buffering by myofilaments that directly disrupts calcium homeostasis (10-12). In this model, one would expect to observe an increased binding affinity for calcium in the troponin complex containing R92Q. 3) R92Q could alter the distribution of positions assumed by tropomyosin along the thin filament, leading to changes in the fraction of bound myosin crossbridges (13). In this model, one would expect to see changes in the equilibrium constants that define the positioning of tropomyosin along the thin filament. The mechanistic differences between these models have important implications for the design of therapeutic strategies.

Here, we set out to identify the initial molecular insult in troponin T caused by the R92Q mutation, and to link the molecular defect to observed derangements in cellular function. To do this, we developed a human R92Q model in gene-edited human induced pluripotent stem cell-derived cardiomyocytes (hiPSC-CMs). We show here that the initial biophysical insult is altered positioning of tropomyosin along the thin filament. The altered positioning of tropomyosin directly affects cellular tension, leading to secondary adaptive changes in calcium homeostasis, gene expression, and electrophysiology. Our results implicate mechanobiological signaling as a primary driver of disease pathogenesis in HCM.

## Results

### Generation of gene-edited stem cell-derived cardiomyocytes

We used CRISPR/Cas9 to generate two independent human induced pluripotent stem cell (hiPSC) lines that are homozygous for the R92Q mutation (Supplementary Fig. S1). Homozygous lines were used to facilitate direct correlation of the molecular insult with alterations in cellular function. Heterozygous lines would better mimic the disease seen in humans but would contain complex mixtures of wild type (WT) and mutant proteins, confounding the correlation of the molecular and cellular results. Both WT and R92Q hiPSCs were derived from the same parent line and are therefore isogenic except for the mutation. We previously showed, by whole exome sequencing of the parent line, that it has no known variants associated with cardiomyopathy (14). Gene-edited hiPSCs have normal karyotypes (Fig. S1B) and are pluripotent, as assessed by immunofluorescence (Supplementary Fig. S2). hiPSCs were differentiated to hiPSC-CMs through temporal modulation of WNT signaling (15, 16), and our efficiency of differentiation using this procedure is >90% (14).

### R92Q hiPSC-CMs generate increased force, power, and contraction speed compared to WT cells

To test whether R92Q hiPSC-CMs show the altered contractility seen in some model systems, we measured the contractility of single hiPSC-CMs using traction force microscopy. hiPSC-CMs were seeded onto rectangular extracellular matrix (ECM) patterns on polyacrylamide hydrogels of physiological stiffness (10 kPa) (14). This patterning on physiological stiffness hydrogels promotes hiPSC-CM maturation and sarcomeric alignment (17). The force, speed of contraction, and power were calculated from the displacement of beads embedded in the hydrogel (18). Data were plotted as cumulative distributions of single cells to account for cell-to-cell variability (14). R92Q hiPSC-CMs generate more force, power, and have a higher contractile speed compared to the WT (Fig. 2).

**Figure 2.**
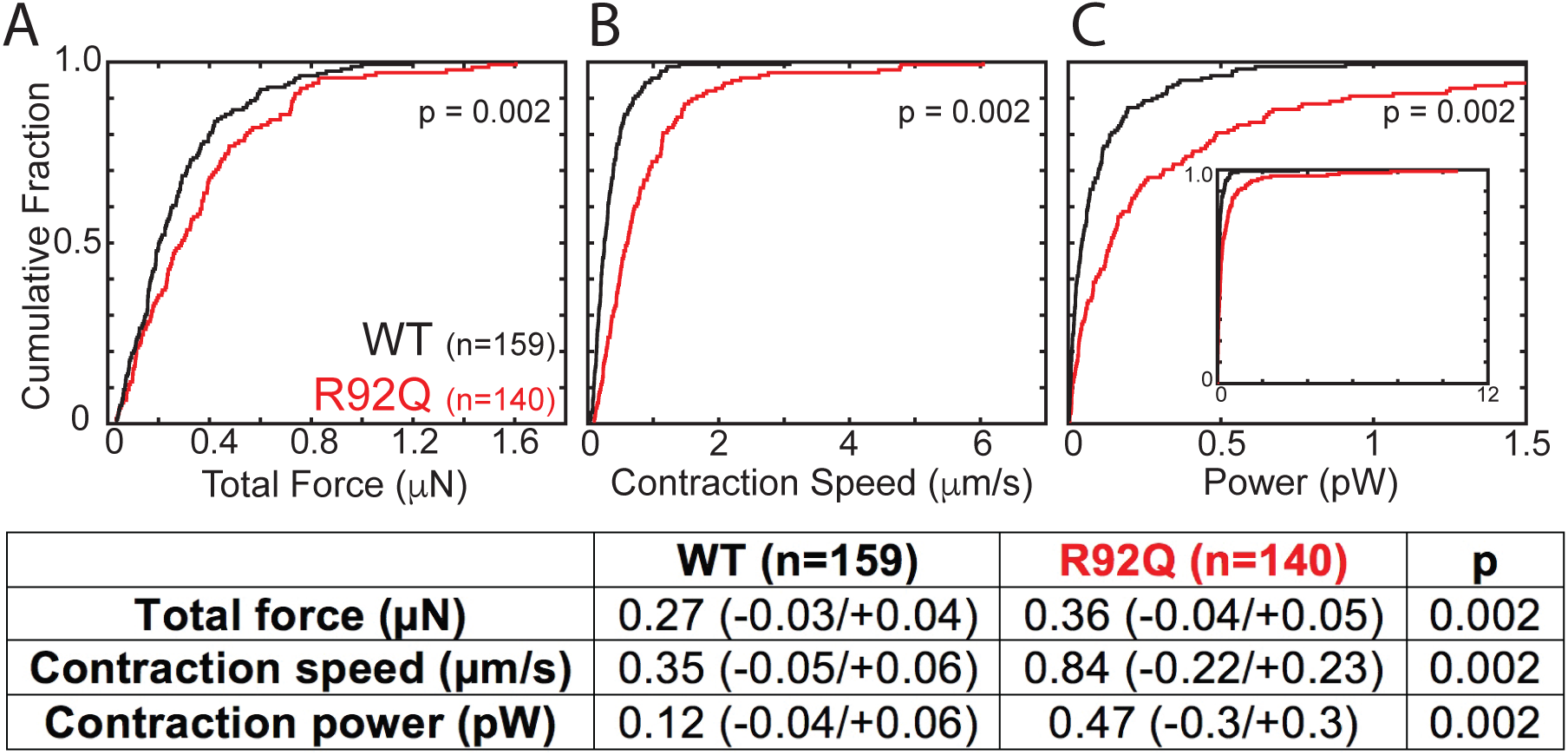
R92Q causes cellular hypercontractility in hiPSC-CMs. Single hiPSC-CMs were seeded on rectangular patterns on 10 kPa hydrogels for traction force microscopy. Cumulative distributions reveal that R92Q hiPSC-CMs have a (A) greater total force, (B) contraction speed, and (C) contraction power compared to the WT. Values from the analysis, 95% confidence intervals, and p-values are listed in the table.

### Intracellular calcium transients are reduced in R92Q cells

Previous studies using R92Q transgenic mice showed altered cardiomyocyte calcium handling (10, 11, 19, 20). To examine calcium dynamics in hiPSC-CMs, cells were patterned onto rectangular ECM patterns on 10 kPa hydrogels and loaded with the ratiometric fluorescent calcium indicator dye, Fura Red. Line scans of the fluorescence of spontaneously beating cells were collected at 1.9 ms intervals. As can be seen, hiPSC-CMs display well-defined calcium transients (Fig. 3A); however, the amplitudes of the transients are lower (p < 0.002) in R92Q (0.56 ± 0.13; n=18), compared to WT (0.84 ± 0.11; n=19), cells. Therefore, despite generating increased force, R92Q hiPSC-CMs show reduced calcium transient amplitudes compared to WT cells.

**Figure 3.**
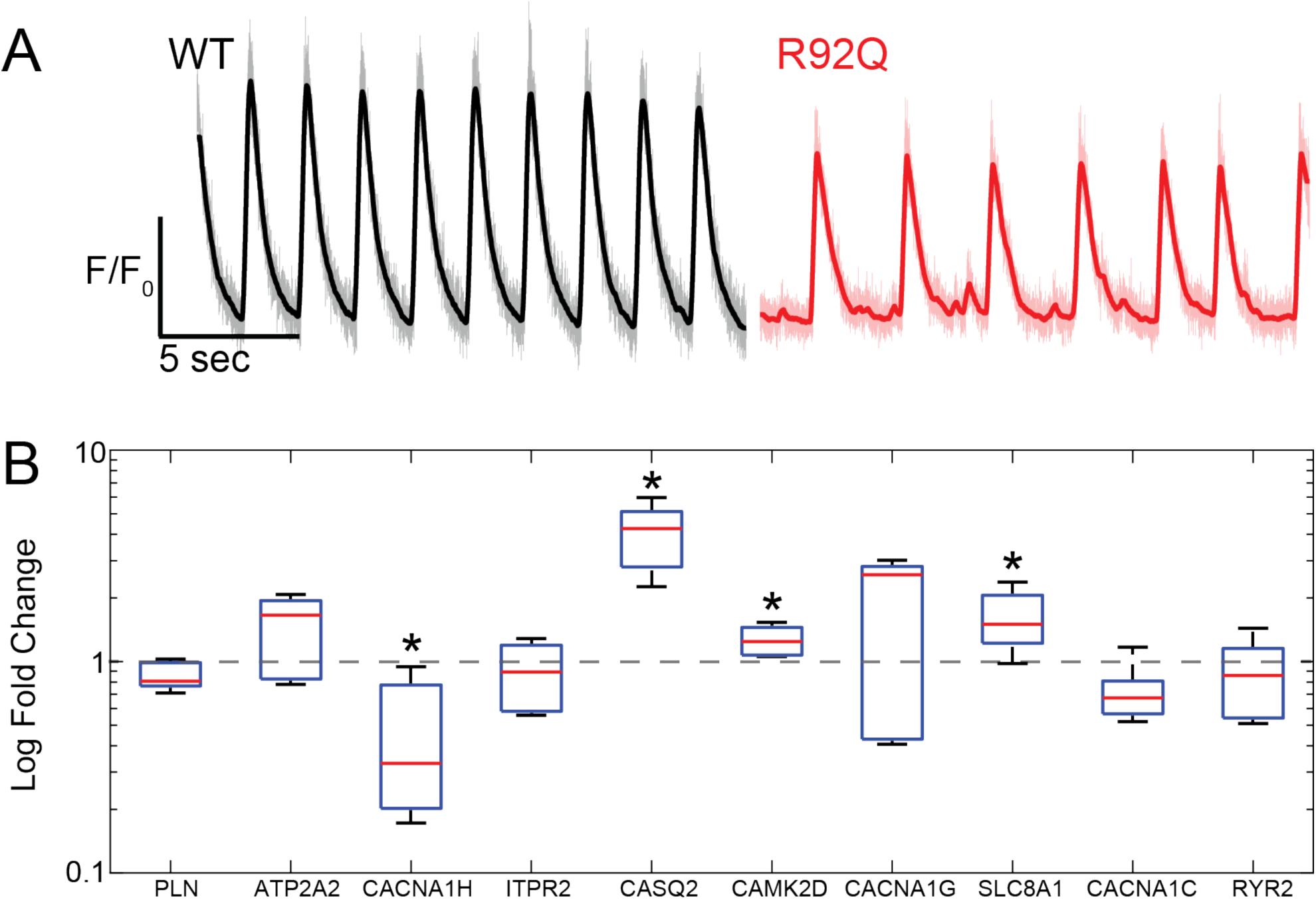
R92Q hiPSC-CMs show altered calcium transients and gene expression. (A) Representative fluorescence traces showing calcium transients. Single hiPSC-CMs were seeded on rectangular patterns on 10 kPa hydrogels and loaded with the ratiometric calcium dye, Fura Red. R92Q hiPSC-CMs calcium transients have lower amplitudes than the WT cells. (B) Expression of key calcium-handling genes measured using qPCR. Data show significant increases in the expression of CASQ2, CAMK2D, and SLC8A1, and a decrease in CACNA1H. ΔCt values are shown in Supplementary Table S2. Statistics were performed on the ΔCt values; however, we show the log-fold change. Red lines show the means, boxes show the quartiles, and error bars show the standard deviations. Data are collected from 3 biological replicates, each of which contained 3 technical replicates. * denotes ΔCt values with p<0.05 compared to the WT.

### R92Q cells show alterations in expression of calcium-handling genes

The observed changes in calcium handling could come from a variety of sources, including changes in transcription, protein expression, and/or post-translational modifications of proteins that regulate calcium homeostasis. To explore a possible role for transcriptional remodeling, we performed qPCR analyses of the expression of transcripts encoded by key genes involved in the regulation of calcium homeostasis in cardiomyocytes. Specifically, we examined the expression levels of transcripts encoding phospholamban (*PLN*), sarcoendoplasmic reticulum calcium-ATPase (*ATPA2*), voltage-gated calcium channel subunits (*CACNA1C, CACNA1G, CACNA1H*), IP3 receptor (*ITPR2*), calsequestrin (*CASQ2*), calcium-calmodulin dependent kinase 2 (C*AMK2D*), sodium-calcium exchanger (*SLC8A1*), and the ryanodine receptor (*RYR2*). We found marked upregulation of *CASQ, CAMK2D*, and *SLC8A1* and downregulation of *CACNA1H* in R92Q, compared with WT, hiPSC-CMs (Fig. 3B, Supplementary Table S2), demonstrating that the expression levels of key genes associated with calcium handling are altered in R92Q hiPSC-CMs.

### R92Q cells show altered action potentials and reduced inward calcium current densities

The observed reductions in the calcium transients observed in spontaneously beating R92Q cells could reflect changes in transmembrane calcium influx. To determine directly if membrane excitability is altered in R92Q cells, we obtained whole-cell current clamp recordings of spontaneous action potentials in WT and R92Q mutant hiPSC-CMs patterned onto rectangular ECM patterns on 10 kPa hydrogels (Fig. 4A-B). Analyses of the data obtained in these experiments revealed that the maximum diastolic potential (the most negative membrane potential achieved between action potentials in spontaneously firing cells) is more depolarized in R92Q hiPSC-CMs than in WT hiPSC-CMs (Fig. 4B). In addition, the frequency of spontaneous action potential firing is higher, upstroke velocities (i.e., the rate of membrane depolarization) are lower, and action potential durations, measured at 50% repolarization (APD_50_), are shorter in R92Q hiPSC-CMs, compared with WT cells (Fig. 4B, Supplementary Table S3).

**Figure 4.**
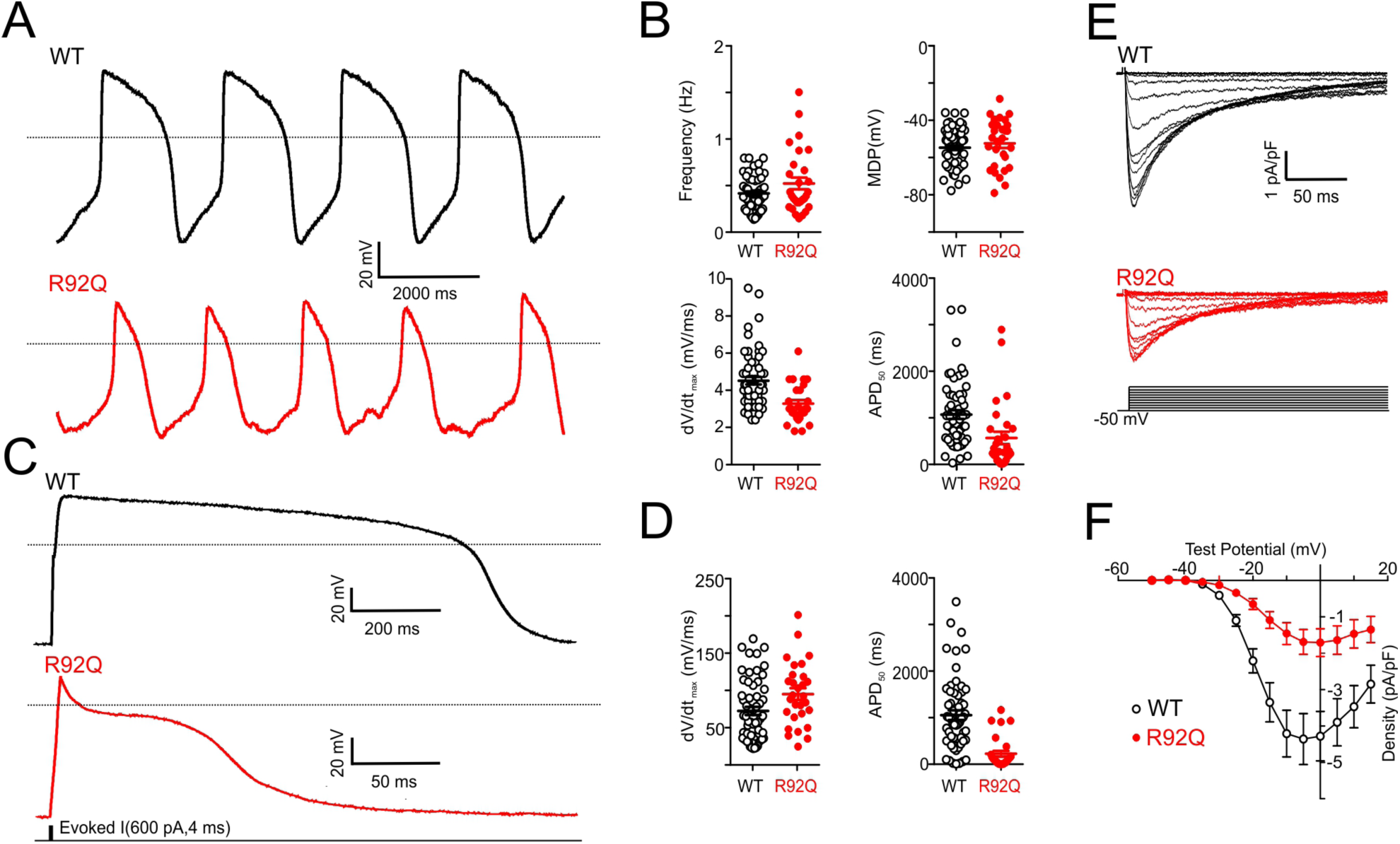
Spontaneous and evoked action potentials are altered in R92Q hiPSC-CMs and I_Ca_ densities are reduced. (A) Representative whole-cell spontaneous action potentials recorded from WT and R92Q hiPSC-CMs are illustrated; dotted black lines indicate 0 mV. (B) Firing frequencies, maximum diastolic potentials (MDP), maximum upstroke velocities (dV/dt_max_) and action potential durations at 50% repolarization (APD_50_), measured in individual WT (n = 58) and R92Q (n = 29) hiPSC-CMs are plotted; mean values are also indicated and are provided in Supplementary Table S3. (C) Representative whole-cell action potential waveforms evoked from a hyperpolarized membrane potential, as described in Material and Methods, in WT and R92Q cells are shown; dotted black lines indicate 0 mV. (D) dV/dt_max_ and APD_50_ values measured in individual WT (n = 58) and R92Q (n = 29) cells are plotted; mean values are also indicated and are provided in Supplementary Table S3. (E) Representative voltage-gated Ca^2+^ current (I_Ca_) waveforms, elicited by voltage steps to test potentials between −40 and +15 mV (in 5 mV increments) from a holding potential (HP) of −50 mV, in WT and R92Q hiPSC-CMs are shown. (F) Mean ± SEM peak I_Ca_ densities in R92Q (n = 12) and WT (n = 15) hiPSC-CMs are plotted as a function of the test potential.

To better understand the mechanism(s) contributing to the reductions in the APD_50_ seen in spontaneously beating R92Q cells, we examined the waveforms of evoked action potentials of hiPSC-CMs hyperpolarized to a membrane potential of −80 mV. Although similar hyperpolarizing currents were required to render R92Q and WT hiPSC-CMs electrically silent and similar currents were required to evoke action potentials in WT and mutant cells, the durations of evoked action potentials are significantly shorter in R92Q, than in WT, cells (Fig. 4C-D).

Additional voltage-clamp experiments were conducted to determine directly if voltage-gated inward calcium current densities were altered in R92Q, compared with WT, cells. With outward potassium currents blocked, we recorded whole-cell voltage-gated calcium currents evoked on membrane depolarization in WT and R92Q hiPSC-CMs. As illustrated in Figure 4E, these experiments revealed that inward calcium current densities are markedly reduced in R92Q, compared to WT hiPSC-CMs (Fig. 4F).

### Determination of the molecular mechanism of R92Q

The cellular studies describe above clearly show changes in cellular mechanics, calcium handling, gene expression, and electrophysiology in R92Q, compared with WT, hiPSC-CMs. However, it is difficult to deconvolve the initial driver of the disease pathogenesis from downstream effects in the inherently complicated cellular context. At the molecular scale, the initial insult that drives the disease pathogenesis is mutation-induced alterations in protein function. Therefore, we set out to determine the molecular mechanism of the R92Q mutation in troponin T.

Troponin T is part of the troponin complex, which, together with tropomyosin, regulates the calcium-dependent interactions between myosin and the thin filament that power force generation in muscle. Biochemical (21) and structural (22) measurements have demonstrated that tropomyosin can lie in three states along the thin filament, termed blocked, closed, and open, and, in addition, that myosin can bind either weakly or strongly to the thin filament when tropomyosin is in the open position (Fig. 1B). In the absence of calcium, tropomyosin lies in the blocked position and inhibits the binding of force-generating actomyosin crossbridges. When calcium binds to troponin C on a thin filament regulatory unit, tropomyosin shifts to the closed position. The tropomyosin can then be pushed into the open position either by thermal fluctuations or myosin binding. Myosin is then able to isomerize into a strong binding state, generating force. The number of strongly bound, force-generating myosin crossbridges at a given calcium concentration determines the amount of force developed.

To examine the molecular effects of the R92Q mutation, WT and R92Q human troponin T were expressed and reconstituted into functional troponin complexes for biochemical and biophysical measurements. All assays were conducted using recombinant human tropomyosin and troponin complex. β-cardiac ventricular cardiac myosin (*MYH7*) and cardiac actin were purified from porcine hearts. The porcine β-cardiac myosin isoform has 97% identity with human β-cardiac myosin (compared to 92% with murine ventricular myosin), and it has very similar biophysical properties, including the kinetics of the myosin ATPase cycle and mechanics measured in the optical trap (23-25).

We examined the effect of the R92Q mutation on thin filament regulation using an *in vitro* motility assay. In this assay, fluorescently labeled reconstituted regulated thin filaments are translocated over a bed of myosin in the presence of ATP and varying concentrations of calcium (26). The speed of translocation was measured as a function of calcium concentration, and normalized data were fitted with the Hill equation, as previously described (27). As can be seen from the data (Fig. 5A), R92Q-regulated thin filaments show a shift towards activation at submaximal, but physiologically relevant, calcium concentrations (pCa_50_ for WT = 6.12 ± 0.02 versus 6.37 ± 0.03 for R92Q; p < 0.001). There is no change in cooperativity, as determined by the Hill coefficient (3.8 ± 0.6 for WT versus 3.4 ± 0.7 for R92Q; p = 0.75).

**Figure 5.**
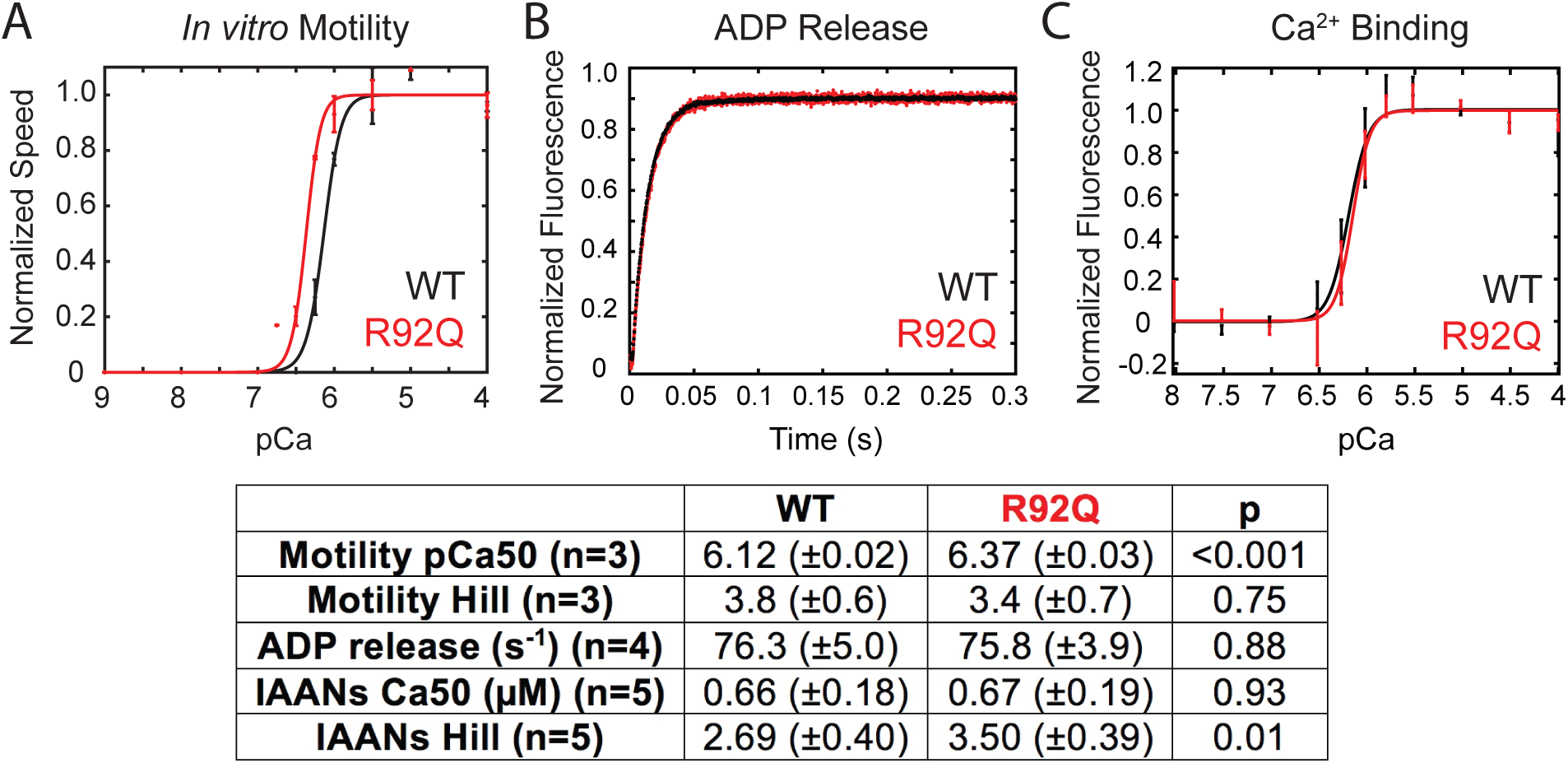
Molecular studies of R92Q demonstrate that R92Q does not change the rate of unloaded actomyosin dissociation or calcium binding affinity to troponin C. (A) *In vitro* motility assays using cardiac myosin and reconstituted regulated thin filaments. The speed of motility was measured as a function of calcium. R92Q shows a shift towards submaximal calcium activation. Error bars are standard deviations from 3 separate experiments. (B) The rate of ADP release from myosin attached to regulated thin filaments was measured using stopped-flow kinetics (fluorescence transients are shown). Myosin bound to ADP and pyrene-labeled regulated thin filaments was rapidly mixed with ATP and the fluorescence increases as myosin dissociates from the thin filament. The rate of actomyosin dissociation was unchanged by the R92Q mutation. (C) The affinity of calcium binding to the troponin complex. IAANS-labeled troponin C was reconstituted into regulated thin filaments. Titrations with increasing calcium were conducted, and there is no difference in calcium binding affinity between the WT and R92Q. Error bars show the standard deviation of 5 experiments. Values derived from fits, standard errors in the fits, and p-values are shown.

### The R92Q mutation does not change myosin detachment kinetics or calcium binding affinity

The shift towards submaximal calcium activation observed for R92Q in the *in vitro* motility assay stems from changes in the function of the troponin-T protein. Given the role of troponin T in regulating calcium-dependent muscle contraction, three models have been proposed to explain the molecular mechanism of the R92Q mutation (Fig. 1B): 1) R92Q could affect the cycling kinetics of myosins that are bound to the thin filament (9). In this model, one would expect to observe a change in the amount of time that myosin remains bound to the thin filament during crossbridge cycling in the mutant. 2) R92Q could increase the calcium affinity of the troponin complex, leading to altered calcium buffering by myofilaments that directly disrupts calcium homeostasis (10-12). In this model, one would expect to observe an increased binding affinity for calcium in the troponin complex containing R92Q. 3) R92Q could alter the distribution of positions assumed by tropomyosin along the thin filament, leading to changes in the fraction of bound myosin crossbridges (13). In this model, one would expect to see changes in the equilibrium constants that define the positioning of tropomyosin along the thin filament. We set out to test these three models.

First, we tested whether the mutation affects the kinetics of myosin detachment from the thin filament by using stopped-flow kinetics to measure the rate of ADP release from actomyosin (i.e., the transition that limits actomyosin dissociation and myosin’s unloaded sliding velocity) (28), as we have done previously (14). We found that the rate of ADP release from myosin bound to regulated thin filaments is not affected by the R92Q mutation (75.8 ± 3.9 s^-1^ for R92Q versus 76.3 ± 5.0 s^-1^ for WT; p = 0.88) (Fig. 5B). Therefore, changes in myosin detachment kinetics cannot explain the shift towards submaximal calcium activation seen in the *in vitro* motility assay.

Next, we measured whether the calcium binding affinity to the troponin complex is affected by the mutation. We used an IAANS-labeled form of troponin C to characterize calcium binding to the troponin complex (29, 30). The fluorescence intensity of this probe changes upon calcium binding to troponin C (29-32). We used it to spectroscopically measure the affinity of calcium binding to regulated thin filaments (Fig. 5C) (29). We saw that the calcium concentration required for half maximal activation, Ca_50_, is not significantly different for the WT (0.66 ± 0.18 μM) and R92Q mutant (0.67 ± 0.19 μM; p = 0.93) proteins. Similar results were seen at 15°C (Fig. 5C) and 20°C (Supplementary Fig. S3). These results demonstrate that changes in the affinity of calcium binding to troponin C cannot explain the shift towards submaximal calcium activation seen in the *in vitro* motility assay (Fig. 5A).

### The initial biophysical insult of R92Q is increased thin filament activation due to repositioning of tropomyosin along the thin filament

To test whether the shift in calcium sensitivity can be explained by a change in the distribution of positions assumed by tropomyosin along the thin filament (Fig. 6A), we measured the equilibrium constants that define the fraction of thin filament regulatory units in each state (21, 33). The equilibrium constant between the blocked and closed states, K_B_, was determined by rapidly mixing fluorescently labeled regulated thin filaments together with myosin and then measuring the rate of myosin binding (seen as quenching of the fluorescence signal) in the presence and absence of calcium (see Materials and Methods for details). At low calcium, when tropomyosin is primarily in the blocked state, the rate of myosin binding to the thin filament is slower than at high calcium, when the blocked state is less populated. The ratio of the rates of binding at low and high calcium were used to calculate K_B_ (Eq. 1, Fig. 6B). As can be seen from the fluorescence transients, the rate of myosin binding to regulated thin filaments is similar for the WT and R92Q mutant proteins at high calcium (pCa 4); however, at low calcium (pCa 9), the rate of binding for the mutant is much faster than for the WT, consistent with lower population of the blocked state. When we calculate K_B_, we see that it is significantly larger in the mutant compared to the WT (1.02 ± 0.26 for R92Q vs. 0.40 ± 0.15 for WT, p=0.003), meaning that the population of the more inhibitory blocked state is reduced while the population of the closed state is increased. The increased K_B_ value means that, at low calcium levels, the thin filament will be more activated in the mutant, consistent with the *in vitro* motility measurements (Fig. 5A).

**Figure 6.**
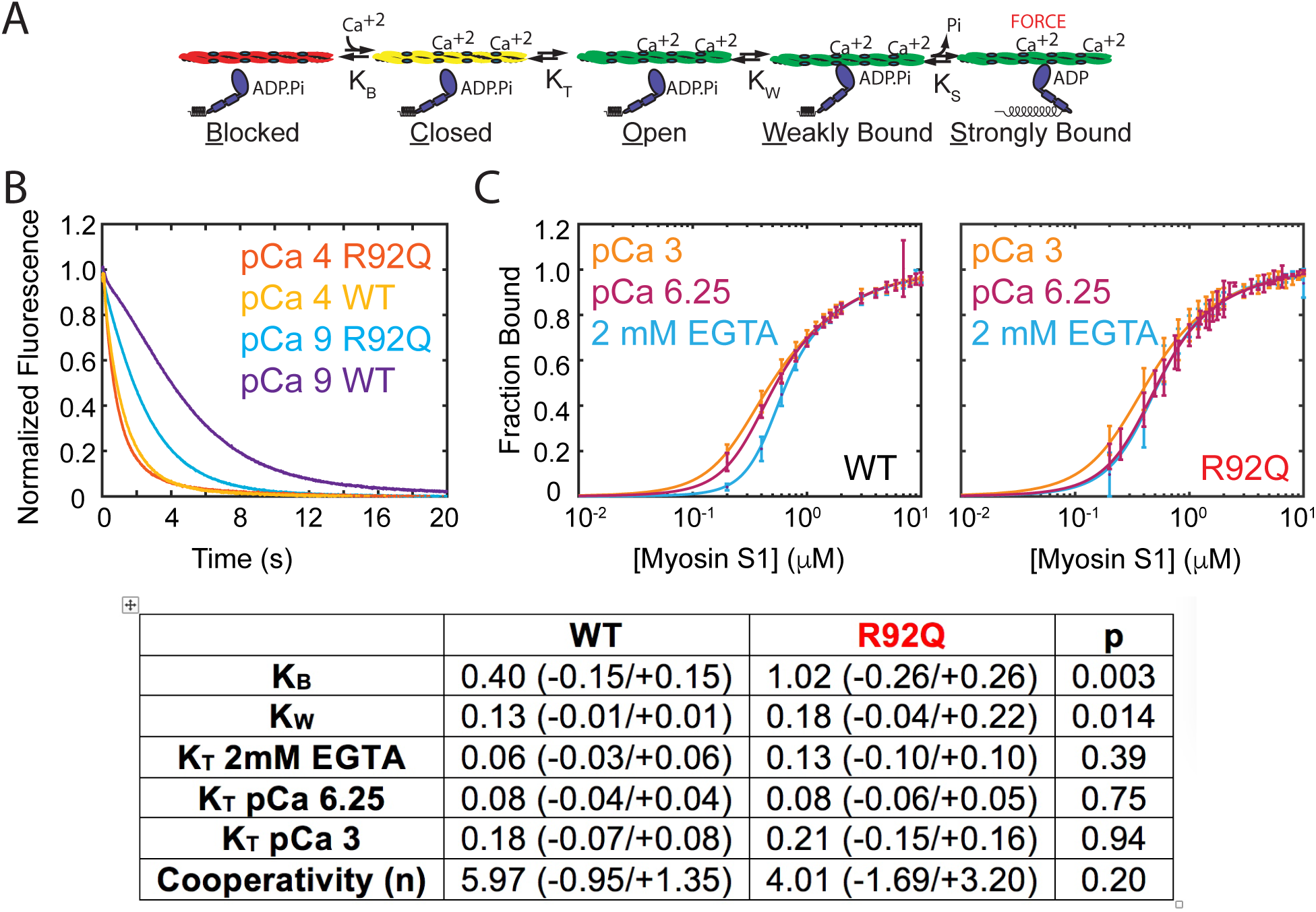
R92Q alters the positioning of tropomyosin along the thin filament. (A) Kinetic scheme for thin filament activation. (B) Measurement of the equilibrium constant K_B_ using stopped-flow kinetics methods (fluorescence transients are shown). Pyrene-labeled regulated thin filaments were rapidly mixed with myosin at high (pCa 4) or low (pCa 9) calcium, and the rate of myosin binding was measured by quenching of the fluorescence. K_B_ is calculated as described in the Methods. The rate of myosin binding is similar for the WT and R92Q at pCa 4, but much faster for R92Q at pCa 9, consistent with destabilization of the blocked state. K_B_ for R92Q is significantly larger than the WT. (C) Measurement of the parameters K_T_ and K_W_ using equilibrium titrations of myosin with regulated thin filaments (see Methods). Fitting reveals no significant differences for R92Q for either K_T_ or K_W_ compared to the WT. Error bars show the standard deviation of 5 experiments. The average value, 95% confidence intervals, and p-values are shown in the table.

Next, we considered whether the mutation affects the equilibrium constant for the transitions between the closed and open states, K_T_, or the equilibrium constant between the open and myosin weakly bound states, K_w_. To do this, we performed titrations of fluorescently labeled regulated thin filaments with increasing concentrations of myosin and measured the quenching of the fluorescence as the myosin binds to the regulated thin filaments (Fig. 6C). The data, analyzed using a modification of the method of McKillop and Geeves (21, 33), show that there are no significant differences in K_T_ between the WT and R92Q (Fig. 6). There is a statistically significant increase in K_w_; however, this is small, and the magnitude is insufficient to explain the shift in the *in vitro* motility assays. This demonstrates that the primary molecular defect in R92Q is partial activation of the thin filament at low calcium levels due to reduced population of the inhibitory blocked state. Based on this result, one would expect increased contractility in the mutant, compared to the WT, during a calcium transient.

### Computational modeling demonstrates that altered tropomyosin positioning with R92Q is sufficient to explain the increase in cellular contractility

To test whether the observed change in K_B_ is sufficient to explain the shift towards submaximal calcium activation seen in the *in vitro* motility assay (Fig. 5A), we used a computational model of thin filament activation developed by Campbell et al (34). In this model, the user inputs a set of equilibrium constants, and the model predicts several parameters, including the force per sarcomere as a function of calcium. When we use the default parameters of the model, but proportionally increase the value of K_B_ to match the fractional change seen in our biochemical experiments, we find that this change alone produces a shift towards submaximal calcium activation similar to the shift observed in the *in vitro* motility experiments (Fig. 7A). This finding validates that the primary effect of the R92Q mutation on motility can be explained by reduced population of the thin filament blocked state.

**Figure 7.**
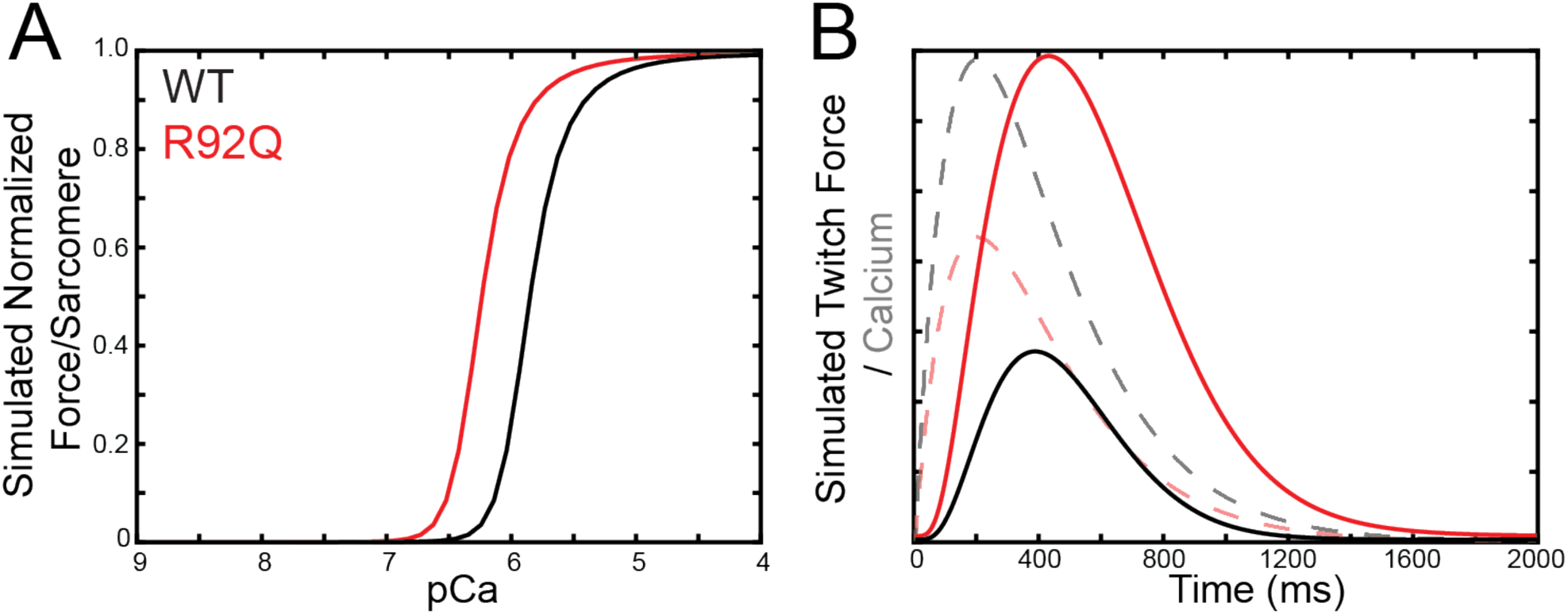
Computational modeling reveals that altered tropomyosin positioning is sufficient to explain the hypercontractility seen with R92Q. (A) Using the computational model developed by (34) and the measured equilibrium constants for thin filament activation, the steady state force per sarcomere was calculated (see Materials and Methods). Changing K_B_ alone is sufficient to reproduce the shift towards submaximal calcium activation seen in the *in vitro* motility experiments (Figure 5A). (B) Using the same model, the equilibrium constants measured *in vitro*, and the calcium transients measured in hiPSC-CMs, the twitch force (solid line) in response to a calcium transient (dashed line) was calculated. Consistent with our cellular measurements, the simulations demonstrate that despite having a reduced calcium transient, R92Q produces a larger force in a twitch due to changes in tropomyosin positioning.

Our data with hiPSC-CMs demonstrate that R92Q has increased force production (Fig. 2A), but reduced calcium transient amplitudes (Fig. 3A). To see whether the reduced population of the blocked state, observed in our molecular studies (Fig. 6A), is sufficient to explain the hypercontractility seen in cells despite the reduction in calcium transient amplitude, we used the same computational model to calculate the expected force per sarcomere in response to a calcium transient. In the modeling, the amplitude of the calcium transient for R92Q was reduced to 67% of the value seen in the WT, as observed in our cellular measurements (Fig. 3A). As above, we proportionally increased K_B_ for the mutant to match the relative difference seen in our biochemical experiments. Consistent with our cellular experiments, the model predicts that the mutant will generate more force in response to a calcium transient than the WT, despite having a smaller amplitude calcium transient (Fig. 7B). Taken together, our molecular experiments demonstrate that the initial molecular insult of the R92Q mutation is decreased population of the thin filament blocked state, leading to increased force generation during a calcium transient.

## Discussion

Here, we elucidated the molecular and cellular consequences of the R92Q mutation in troponin T that causes HCM, R92Q. We show that the initial molecular insult that drives disease pathogenesis is increased thin filament activation at physiologically relevant micromolar calcium levels due to destabilization of the blocked state of tropomyosin. We demonstrate computationally and experimentally that this increased activation directly causes increased mechanical force produced by hiPSC-CMs. We show that this initial insult of altered mechanical forces leads to downstream changes in the expression of genes associated with calcium handling, altered calcium transients, and alterations in cellular electrophysiology. Taken together, our results highlight the role of mechanobiology in driving the early disease pathogenesis.

### Defining the primary molecular driver of the disease pathogenesis

Previous *in vivo* and *in vitro* cellular studies have demonstrated that the R92Q mutant protein is expressed and properly integrated into sarcomeres, suggesting that the driver of the disease is changes in protein biochemistry and biophysics, rather than haploinsufficiency (4, 7, 8, 35). To better understand these changes in protein function, we conducted *in vitro* motility assays which demonstrated that R92Q causes a shift towards submaximal calcium activation (Fig. 5A). This finding is consistent with some (6, 9, 35-39), but not all (5, 7), previous measurements in muscle fibers and in biochemical assays using non-cardiac muscle protein isoforms. The shift towards submaximal calcium activation could potentially come from changes in actomyosin dissociation kinetics, the affinity of calcium binding to troponin C, and/or the positioning of tropomyosin along the thin filament (Fig. 1B).

Our biophysical studies clearly demonstrate that the mutation does not affect the binding of calcium to troponin C or the kinetics of actomyosin dissociation in the absence of load (Fig. 5). The results show, however, that the mutation causes a pronounced increase in the equilibrium constant between the blocked and closed states, K_B_ (Fig. 6B). This change would favor the closed state over the blocked state, effectively lowering the energy barrier required for thin filament activation at physiologically relevant (pCa 5-7 range (40)) calcium concentrations. Our computational modeling (Fig. 7) demonstrates that the observed change in this equilibrium constant is sufficient to explain the shift towards submaximal calcium activation seen in our *in vitro* motility measurements (Fig. 5A). Our data support a model in which the initial insult that drives disease pathogenesis is altered positioning of tropomyosin along the thin filament, with a greater fraction of regulatory units in the closed, than in the more inhibitory blocked, position at low calcium. This shift would lower the energy barrier for activation of the thin filament, leading to submaximal calcium activation.

The R92Q mutation has been studied in many model systems, including quail myotubes (7), transfected rat cardiomyocytes (5), skinned rabbit muscle fibers (6), transgenic mice (8), and transfected cat cardiomyocytes (4). While these studies have greatly advanced our understanding of the mutation, they have also shown that the effects of the mutant protein present differently depending on the model system used. In addition, previous work using transgenic mice demonstrated that disease presentation varies depending on whether proteins with biophysical properties similar to human isoforms are used (9). The use of all cardiac proteins with biochemical and biophysical properties similar to human proteins is especially important for studies of thin filament mutations, since the activation of the thin filament depends on both myosin and calcium binding (Fig. 1B). In our molecular studies, we used human cardiac troponin and tropomyosin and porcine cardiac myosin and actin. Porcine cardiac myosin (*MYH7*) is 97% identical to the human protein, and displays biochemical kinetics, mechanical step sizes, and load-dependent kinetics that are indistinguishable from the human isoform (23-25). Therefore, we believe that our model system reliably mimics the molecular phenotype in humans.

Interestingly, the R92 residue is in the region of troponin T that interacts with tropomyosin, near where two tropomyosin molecules overlap in a head-to-tail fashion (41). Two other HCM-causing mutations have been identified at the R92 site, R92W and R92L, and this has led to the suggestion that it is a hotspot for HCM mutations. To date, there are no atomic-resolution structures of this region of the thin filament. Structural studies, however, have shown that troponin T plays a role in stabilizing the blocked state in the absence of calcium (42). In addition, molecular dynamics simulations have shown that mutations in the R92 region can lead to changes in the distance between troponin T and tropomyosin (43) and biochemical experiments have shown that the R92L mutation decreases the affinity of troponin for tropomyosin (44). We speculate that the R92Q mutation has a similar effect on the interactions between troponin T and the thin filament, leading to destabilization of the blocked state.

It has previously been proposed that R92Q causes an increase in calcium affinity for the troponin complex on the thin filament which would affect the buffering of calcium by myofilaments, leading to disrupted calcium homeostasis (10, 12). While our cellular data reveal disrupted calcium homeostasis, our molecular work shows no change in the affinity of calcium for R92Q troponin, demonstrating that this change in calcium homeostasis is a downstream consequence of the primary molecular insult. This result is consistent with work from the Molkentin lab (45), which showed that the development of HCM correlates with changes in tension, rather than calcium handling.

Recent studies of HCM-causing mutations in thick filament proteins, including β-cardiac myosin (*MYH7*), myosin binding protein C (*MYBPC3*), myosin regulatory light chain, and myosin essential light chain, have demonstrated that many of these mutations disrupt the autoinhibited super relaxed state of myosin, leading to the recruitment of more crossbridges and thus hypercontractility (46-51). It has been proposed that increased crossbridge recruitment correlates with the hyperdynamic cardiac function seen in HCM (48). Our studies with R92Q, a thin filament mutation, demonstrate a similar net effect of increased crossbridge recruitment at physiologically relevant calcium levels, suggesting altered recruitment of crossbridges in HCM as a common theme for both thin and thick filament mutations.

### Connecting the molecular and cellular phenotypes in R92Q

Our data clearly demonstrate that the primary molecular driver of early disease pathogenesis is altered positioning of tropomyosin along the thin filament. At the cellular level, R92Q shows both an increase in cellular force production (Fig. 2A) and a reduction in the amplitude of the calcium transient (Fig. 3A). These seemingly conflicting findings can be reconciled by our computational modeling (Fig. 7B), which reveals that the shift towards thin filament activation at submaximal calcium leads to cellular hypercontractility, despite the reduction in the amplitude of the calcium transient. The hypercontractile effects of this shift are relevant at physiological (micromolar) concentrations of calcium (52). Importantly, our results demonstrate that the cellular hypercontractility can be explained by our molecular mechanism.

At the cellular level, we see disrupted calcium homeostasis with R92Q, which is a downstream consequence of the primary hypercontractile phenotype. Calcium homeostasis in the myocardium is a complicated process which depends on many factors, including gene expression and post-translational modifications of signaling and contractile proteins (52). While a complete dissection of this mechanism is beyond the scope of the current study, our work provides insights into potential transcriptional mechanisms. We observed changes in the expression of several genes involved in calcium handling (Fig. 3B), including calsequestrin (*CASQ2*), calcium-calmodulin kinase (*CAMK2D*), the sodium-calcium exchanger (*SLC8A1*), and a voltage-gated calcium channel subunit (*CACNA1H*). Interestingly, overexpression of *CASQ* or *CAMK2D* in transgenic mice drives the development of heart failure and arrhythmogenesis (53, 54). We recognize that changes in transcript expression do not always correlate with protein function. Regardless, our data demonstrate that altered mechanics at the molecular level can drive changes in cellular gene expression, showing a mechanobiological link between these processes in HCM.

At a functional level, our single-cell electrophysiological experiments reveal that action potential durations are shorter in R92Q, compared with WT cells, due in part to reduced inward L-type calcium current densities (Fig. 4). These changes would be expected to be arrhythmogenic and could contribute to the increased incidence of arrhythmias and sudden cardiac death in individuals harboring the R92Q mutation. We observe normal expression levels of the transcripts encoding L-type channel subunits, and therefore, the reduced current density could be due to alterations in signaling pathways and/or post-translational modifications of channel subunit proteins. The reduction in inward calcium current densities observed in our electrophysiological measurements (Fig. 4) would be expected to reduce calcium-induced calcium release from intracellular calcium stores, potentially contributing to the observed reductions in calcium transient amplitudes (Fig. 3A).

Our results clearly show that molecular hypercontractility drives downstream changes in cellular calcium handling and electrophysiology in single hiPSC-CMs. We propose that mutation-induced changes in cellular tension alter mechanosensitive signaling pathways in cardiomyocytes (55). Consistent with this idea, recent work from our lab demonstrated that a dilated cardiomyopathy mutation in troponin T, ΔK210, affects molecular mechanosensing, which helps to drive the disease progression (14). Such a mechanism is also consistent with the model of Davis et al., who proposed that alterations in cellular tension correlate with the hypertrophic response (45). In fact, increases in cardiac tension stemming from external sources such as hypertensive disease and aortic stenosis can promote pathological hypertrophy. Deciphering the specific mediators of mechanobiological pathways in cardiomyocytes is an active field of research (55-58). In the broader context of the myocardium, hypercontractility of cardiomyocytes can impose aberrant stretch on fibroblasts, activating the transition to myofibroblasts (55). Such a mechanism could contribute to the diffuse myocardial fibrosis frequently seen with HCM.

### Implications for modeling and treating HCM

The goal of our study was to connect the initial molecular insult with the early disease pathogenesis in human cells. We therefore used genome-edited hiPSC-CMs, which are excellent tools for dissecting early disease pathogenesis (59, 60). These experiments were conducted using isogenic cells, making it easier to understand the direct consequences of the point mutation on a controlled genetic background (61). The results obtained demonstrate that these hiPSC-CMs recapitulate important aspects of HCM-induced changes in contractility (8, 9, 37), altered electrophysiology (10, 20), and calcium dysfunction (10, 11, 19) seen in other model systems. While our hiPSC-CM model recapitulates some aspects of the early disease pathogenesis, it cannot fully capture the clinical phenotype seen in patients for several reasons. First, hiPSC-CMs are developmentally immature, and they lack many of the physiological cues present inpatients (59, 60). As such, they do not capture some aspects of clinical HCM, including fibrosis, tissue hypertrophy, and ventricular arrhythmias. Moreover, while patients are typically heterozygous for the R92Q mutation, our studies were conducted using homozygous cell lines to facilitate connecting the molecular and cellular phenotypes. Work in transgenic mice has shown that disease phenotypes vary with mutant gene dosage (8, 62) and that the homozygous mutation is embryonic lethal. Therefore, care should be taken when extrapolating from these studies to the clinical phenotype.

Limitations aside, hiPSC-CMs are a unique tool to study the connection between the initial molecular insult and the early disease pathogenesis in human cells. Our identification of altered cellular mechanics and downstream mechanobiological signaling pathways as key drivers of the disease pathogenesis has important implications for treatment. There is currently an outstanding need to develop new therapeutics to treat HCM. The current therapeutic regimen is the use of agents to prevent further myocardial remodeling, and in extreme cases, myectomy or cardiac ablation. Our findings suggest that approaches which target mechanobiological signaling pathways in cardiomyocytes could be useful in the treatment of HCM.

Recently, there was a report of an HCM mutation in α-actinin that causes prolongation of the action potential due to an increase in the calcium current density (63). In this case, the patient was successfully treated with the L-type calcium channel blocker, diltiazem. In R92Q, we observed a reduction in the calcium current density (Fig. 4), and therefore, a different therapeutic would be necessary. These differences between cellular phenotypes in these two HCM mutations highlights the need to understand the underlying changes in molecular and cellular function, and it demonstrates the need to consider a personalized medicine approach for HCM.

### Conclusions

The results here demonstrate that the initial insult of the R92Q mutation in troponin T is molecular and cellular hypercontractility at physiologically relevant calcium concentrations, which leads to alterations in mechanobiological signaling pathways that regulate calcium homeostasis, gene expression, and cellular electrophysiology. Taken together, the data presented suggest that these mechanobiological adaptations play a central role in the early disease pathogenesis, and they suggest that targeting these pathways could open new avenues for treating this devastating class of diseases.

## Acknowledgements

The authors would like to thank Jonathan Davis for the troponin-C^T53C^ plasmid. The authors acknowledge financial support from Washington University in St. Louis and the Institute of Materials Science and Engineering for the use of instruments and for staff assistance. The authors would also like to acknowledge the financial support provided by the National Institutes of Health (R01 HL141086 to M.J.G., R01 HL034161 and R01 HL142520 to J.M.N.), the March of Dimes Foundation (FY18-BOC-430198 to M.J.G.), the Children’s Discovery Institute of Washington University and St. Louis Children’s Hospital (PM-LI-2019-829 M.J.G.), and the Washington University Center for Cellular Imaging (WUCCI) (CDI-CORE-2015-505 to M.J.G.). S.R.C. was supported through an institutional training grant (T32 EB018266).

## Conflict of interest statement

All experiments were conducted in the absence of any commercial or financial relationships that could be construed as potential conflicts of interest.

## Author contributions

S.R.C. purified proteins and performed and analyzed the stopped flow and fluorescence experiments. P.E.C. performed and analyzed the traction force microscopy experiments with the stem cell derived cardiomyocytes. W.W. performed and analyzed electrophysiological experiments. L.G. purified proteins, implemented the cell-based assays, performed and analyzed experiments with stem cell derived cardiomyocytes, performed qPCR measurements, and performed calcium imaging experiments. W.T.S. designed tools for microcontact printing. P.A. performed *in vitro* motility assays. J.M.N. oversaw the electrophysiological experiments and analyzed data. M.J.G. oversaw the project, performed simulations, generated mutant proteins, implemented biochemical assays, analyzed data, and drafted the manuscript. All authors contributed to the writing and/or editing of the manuscript.

## Methods

### Protein modification and purification

Cardiac myosin and actin were purified from cryoground porcine ventricles (Pelfreez) as previously described (14). S1 myosin was prepared by chymotrypsin digestion as previously described (14). Recombinant human cardiac tropomyosin, troponin I, troponin T, and troponin C were expressed in *E. coli* and purified from BL21-CodonPlus cells (Agilent) as described previously (14). Purified tropomyosin was reduced in 50 mM DTT at 56°C for 5 minutes and ultracentrifuged to remove aggregates immediately before being used in each assay. The R92Q mutation was introduced into troponin T using QuikChange Site-Directed Mutagenesis (Agilent) and the presence of the mutation was verified by sequencing.

For the studies of calcium binding, we used IAANS (6-((4-((2-iodoacetyl)amino)phenyl)amino)-2-naphthalenesulfonic acid)-labeled troponin C (32). IAANS was custom synthesized by Toronto Research Chemicals. Troponin C^T53C^ was labeled with five-fold molar excess IAANS dye overnight, and the reaction was quenched with DTT. Excess dye was dialyzed out with 4 dialysis buffer changes of 1 mM DTT, 0.01% NaN_3_, 50 μM CaCl_2_, 1 mM MgCl_2_, 3 M Urea, 1 M KCl, 5 mg/L TPCK, 5 mg/L TLCK, 0.3 mM PMSF (32). The IAANS-labeled troponin C^T53C^ was then purified over a MonoQ column and complexed with the troponin T and I as done previously (14).

### *In vitro* motility assays

*In vitro* motility assays were conducted using thin filaments containing R92Q troponin T as previously described (14). Data for WT troponin T are from (14). Briefly, enzymatically inactive full-length porcine cardiac myosin was removed by cosedimentation with phalloidin-stabilized F-actin in the presence of ATP. Flow cells were loaded with 1 volume (50 μL) of 200 nM myosin, 2 volumes of 1 mg/mL BSA, 1 volume of 1 μM F-actin, 2 volumes of KMg25 (25 mM KCl, 4 mM MgCl_2_,1 mM EGTA, 1 mM DTT, 60 mM MOPS pH 7.0) + 1 mM MgATP, 4 volumes of KMg25, and 1 volume of 40 nM rhodamine-phalloidin-labeled thin filaments. After loading 2 volumes of activation buffer (KMg25 with 4 mM MgATP, 1 mg/mL glucose, 192 U/mL glucose oxidase, 48 μg/mL catalase, 2 μM troponin and tropomyosin, 0.5% methyl cellulose), flow cells were imaged for 20 frames. Individual motile filaments were manually tracked using the MTrackJ plugin in Fiji ImageJ (64), and each point shows the average and standard deviation of the speed from 3 separate experiments.

### Stopped-flow transient kinetic measurement of K_B_ and ADP release

An SX-20 stopped flow apparatus (Applied Photophysics) was used. K_B_ was determined as previously described (14, 33). WT data are from (14). At both low (pCa 9) and high calcium (pCa 4), 5 μM phalloidin-stabilized pyrene actin, 2 μM tropomyosin, 2 μM troponin, and 0.04 U/mL apyrase were rapidly mixed with 0.5 μM S1 myosin and 0.04 U/mL apyrase. Performed at 20°C, each experiment was the average of at least 3 separate mixes and the data were fit by a single exponential curve. K_B_ was calculated from: 

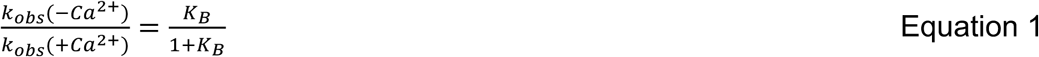

The reported K_B_ is the average of at least three different experiments. The p-value was calculated from a 2-tailed Student’s t-test.

The rate of ADP release from myosin bound to regulated thin filaments (20°C) was measured as previously described. (14) The average and standard deviation of the rate of at least four experiments was calculated and the p-value was derived using a two-tailed Student’s t-test.

### Fluorescence titrations to measure K_W_, K_T_, and n

A SX-20 stopped flow fluorometer was used for all fluorescence titrations. The values of K_W_, K_T_, and n (the cooperativity) were determined for R92Q and WT using fluorescence titrations as previously described (14, 33). The WT data is from (14). Our MATLAB-based computational tool was used for hypothesis testing and uncertainty estimation, as previously described (33).

### Measurement of calcium binding to troponin C

The calcium affinity for the troponin complex (Tn^IAANS^) was determined by titrating regulated thin filaments with increasing calcium concentrations and measuring the change in fluorescence in IAANS-labeled troponin C upon calcium binding (32). Tn^IAANS^ was excited at 330 nm and fluorescence emission was detected using a 395 nm long-pass filter. 0.15 μM Tn^IAANS^ complex, 0.45 μM tropomyosin, and 2 μM actin were mixed with increasing concentrations of calcium in 10 mM MOPS pH 7.0, 150 mM KCl, 3 mM MgCl_2_, and 1 mM DTT. Each buffer was balanced to give the desired free calcium, free magnesium, and ionic strengths using MaxChelator (65). The solution was allowed to equilibrate for 1 minute after mixing with constant stirring before the fluorescence intensity was measured. The titration curve was fit by the logistic sigmoid function, which is mathematically equivalent to the Hill equation: 

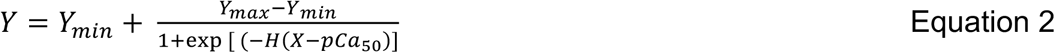

where Y_max_ and Y_min_ are the maximum and minimum IAANS fluorescence, X is the negative logarithm of [Ca^2+^]_free_, pCa_50_ is the negative log of the concentration of free calcium producing half-maximal fluorescence, and H is the cooperativity (proportional to the Hill coefficient) (66). Titrations were performed at both 15°C (Fig. 5C) and 20°C (Supplementary Fig. S3) to facilitate comparison with previous measurements using different proteins (29).

### Computational modeling of sarcomeric contractility

To simulate the effects of the experimentally determined changes in equilibrium constants on force production, we used the computational model developed by Campbell et al. (34) based on McKillop and Geeves (21), as we have done previously (14). Briefly, in this model, 9 sarcomeres are simulated, where the equilibrium constants between states and a coupling constant describing cooperativity define the probability of switching between biochemical states. The steady-state force is calculated from the equilibrium distribution of states at a given calcium concentration. Our biochemical experiments demonstrated that the primary change at the molecular scale with the mutation is an increase in K_B_, such that K_B_ (R92Q) = 2.56 * K_B_ (WT). To simulate the WT, we used the default model parameters. To simulate the mutant, we decreased the reverse rate constant that defines K_B_, so that K_B_ (R92Q) = 2.56 * K_B_ (WT). To simulate the force per sarcomere in response to a calcium transient for the WT, we used the default calcium transient. To simulate the response of R92Q, we changed K_B_ as described above and we reduced the amplitude of the default calcium transient to 67% of its value to match our measurements in hiPSC-CMs.

### Stem cell line derivation

R92Q stem cells were derived and the quality control was performed using procedures described in depth previously (14). Briefly, the parent human BJ fibroblast stem cell line (BJFF.6, ATCC) was reprogrammed to stem cells by the Genome Engineering and iPSC Center (GEiC) at Washington University in St. Louis. Two independent isogenic stem cell lines with the R92Q hTNNT2 point mutation were also generated at the GEiC using the CRISPR/Cas9 system (67). The oligo used to generate the gRNA was CCTTCTCCATGCGCTTCCGGNGG and the mutation was introduced by homology directed repair. This gRNA was selected to minimize off target effects. The presence of the homozygous mutation was verified by sequencing. Karyotype (G-banding) analysis was performed by Cell Line Genetics (Supplementary Fig. S1). Mycoplasma testing and immunofluorescence staining for pluripotency markers were performed by the GEiC.

### Stem cell and hiPSC-CMs culture

Stem cell culture and differentiation to hiPSC-CMs were done as previously described (14). Briefly, stem cells were maintained in feeder-free culture. To differentiate the stem cells to hiPSC-CMs, we used small-molecule manipulation of WNT signaling (15, 16). hiPSC-CMs were enriched using metabolic selection (68). All functional experiments were conducted at least 30 days after the initiation of differentiation. Experiments were conducted using 2 independently derived cell lines for the R92Q mutant. All experiments were repeated using at least two independent differentiations.

### Microcontact patterning of hiPSC-CMs on glass and hydrogels

Fabrication of rectangular (7:1 aspect ratio) PDMS stamps for micropatterning of hiPSC-CMs on both glass and 10 kPa hydrogels was done as previously described (14, 17). Cells were patterned onto 10 kPa polyacrylamide hydrogels containing stamped Geltrex (Thermo Fisher) in rectangular patterns as in (14).

### Traction force microscopy

Traction force microscopy was conducted on 10 kPa hydrogels as previously described (14) and analyzed using the computational tool developed in (18). Data were analyzed and 95% confidence intervals of the mean were calculated as described previously (14, 33).

### Measurement of calcium transients in live cells

Live-cell imaging was conducted using the ratiometric fluorescent calcium indicator dye Fura Red AM (Thermo Fisher). The use of a ratiometric dye is important since the mutation could affect the uptake of dye into the cells, and the ratiometric dye normalizes the calcium-induced changes in fluorescence to the total amount of dye taken up by the cell. hiPSC-CMs were patterned on hydrogels as described above. After 5-7 days on the patterns, the cells were loaded with 10 μM Fura Red AM dye and 0.01% Pluronic F-127 (Invitrogen/ThermoFisher) in RPMI-B27 with insulin media for 20 min at room temperature. The cells were washed twice and incubated with Tyrode’s solution (1.8 mM CaCl2, 135 mM NaCl, 4 mM KCl, 1 mM MgCl2, 5 mM glucose and 10 mM HEPES, pH 7) for 15-20 minutes at 37°C to allow de-esterification of the dye. Calcium transients were recorded with a Nikon A1Rsi confocal microscope in line scan mode using a 40X objective and the Ex2Em1 microscope setting. Fura Red AM loaded cells were excited at both 405 nm and 488 nm, and the emission fluorescence signal was collected at 595nm. Line scans were acquired at a sampling rate of 512 pixels x 1.9 ms per line (total 10,000 lines per recording). Each cell was recorded along with a line scan of the background fluorescence outside the cell area.

### Analysis of calcium transients

The calcium transient fluorescence counts were converted to ratios using Fiji software (64). The averaged background fluorescence was subtracted from each recording and a montage was created from the image stacks. The ratio of fluorescence at 405 nm / 488 nm was then calculated in Excel. The resulting ratiometric calcium fluorescence traces were then analyzed using a custom MATLAB script to calculate the amplitude of the calcium transient. Traces with fewer than 3 peaks were not analyzed. Briefly, the data were smoothed over a 100-point sliding window using a Savitsky-Golay filter. The locations of peaks and minima in the fluorescence signal were determined using a peak-finding algorithm. Statistical significance was tested using a 2-tailed Student’s t-test.

### Measurement of the expression of transcripts encoding key calcium-handling proteins

hiPSC-CMs were grown on Matrigel-coated (Corning) 10 kPa PrimeCoat elastic substrate culture dishes (35 mm) (ExCellness Biotech SA, Lausanne, Switzerland) for 10 days. Total RNA was isolated using RNeasy Mini Kit (Qiagen) with on-column DNase I treatment according to the manufacturer’s instructions. cDNA was generated using iScript Reverse Transcription Supermix (Biorad) according to the manufacturer’s instructions. qPCR reactions were performed in triplicate using iTaq Universal SYBRGreen Supermix (Biorad) and using the ViiA 7 System (Applied Biosystems). Primers for all genes were obtained from IDT PrimeTime qPCR Primers. Primer product numbers from IDT are listed in Supplementary Table S1. Three separate biological samples were evaluated for both WT and R92Q homozygous hiPSC-CMs. The relative levels of mRNA were calculated using the comparative threshold cycle (ΔCt) method (69). GAPDH and HPRT1 were used as endogenous controls, and Rox dye present in the master mix was used to normalize background fluorescence. ΔCt values are plotted in Supplementary Table S2. The statistical significance of differences in ΔCt values was evaluated using a two-tailed Student t-test.

### Cellular electrophysiological studies

Whole-cell current- and voltage-clamp recordings were obtained at room temperature (22∼24°C) from hiPSC-CMs plated on hydrogel-coated coverslips using a Dagan 3900A (Dagan Corporation) amplifier interfaced to a Digidata 1332A A/D converter (Axon) and the pClamp 10.3 software (Axon). For current-clamp recordings, recording pipettes contained 135 mM KCl, 5 mM K_2_ATP, 10 mM EGTA, 10 mM HEPES and 5 mM glucose (pH 7.2; 310 mOsm). The bath solution contained 136 mM NaCl, 4 mM KCl, 2 mM MgCl_2_, 1 mM CaCl_2_, 10 mM HEPES and 10 mM glucose (pH 7.4; 300 mOsm). For recordings of voltage-gated Ca^2+^ currents (I_Ca_), pipettes contained 5 mM NaCl, 90 mM Cs CH_3_SO_3_, 20 mM CsCl, 4 mM MgATP, 0.4 mM Tris-GTP, 10 mM EGTA, 10 mM HEPES and 3 mM CaCl_2_ (pH 7.2; 310 mOsm), and the bath solution contained 20 mM NaCl, 110 mM TEA-Cl, 10 mM CsCl, 1 mM MgCl_2_, 1 mM CaCl_2_, 10 mM HEPES and 10 mM glucose (pH 7.4; 300 mOsm). In all experiments, pipette resistances were 2-3 MΩ.

Electrophysiological data were acquired at 10 or 100 kHz, and signals were low pass filtered at 5 kHz before digitization and storage. After the formation of a gigaohm seal (>1GΩ) and establishment of the whole-cell configuration, brief (10 ms) ± 5 mV voltage steps from a holding potential (HP) of −70 mV were presented to allow measurements of whole-cell membrane capacitances (C_m_), input resistances (R_in_), and series resistances (R_s_). Mean ± SEM C_m_ values were 32 ± 2 pF and 47 ± 2 pF (*P* < 0.001) in WT (n = 96) and R92Q (n = 67), hiPSC-CMs, respectively; mean ± SEM R_in_ values were 1665 ± 125 MΩ and 1551 ± 162 MΩ (*P* > 0.05) in WT (n = 96) and R92Q (n = 67), hiPSC-CMs, respectively. Whole-cell C_m_ and R_s_ were electronically compensated by 85%. Voltage errors resulting from the uncompensated R_s_ were always <2 mV and were not corrected. Leak currents were always <50 pA and also were not corrected.

In current-clamp recordings, spontaneous action potentials were recorded on establishing the whole-cell configuration. To record evoked action potentials, small (−10 ∼ −100 pA) current injections were made to hyperpolarize the membrane potential to −80 mV and to stop spontaneous firing. Individual action potentials were then evoked by brief (4 ms) depolarizing current (600 pA) injections. In voltage-clamp experiments, whole-cell I_Ca_, evoked in response to depolarizing (300 ms) voltage steps to test potentials between −45 and +15 mV (in 5 mV increments at 1 s intervals) from a holding potential of −50 mV, were recorded.

### Analysis of electrophysiological data

Electrophysiological data were compiled and analyzed using Clampfit 10.3 (Axon) and GraphPad (Prism). C_m_ values were determined by integration of the capacitive transients recorded during ± 5 mV voltage steps from −70 mV. Current amplitudes in each cell were normalized to the C_m_ and current densities are reported (pA/pF). All data are presented as means ± SEM. The statistical significance of observed differences between WT and R92Q hiPSC-CMs was evaluated using two-tailed Student’s t-test or two-way ANOVA; p-values are presented in Supplementary Table S3.

---

**Supplementary Figure S1.**
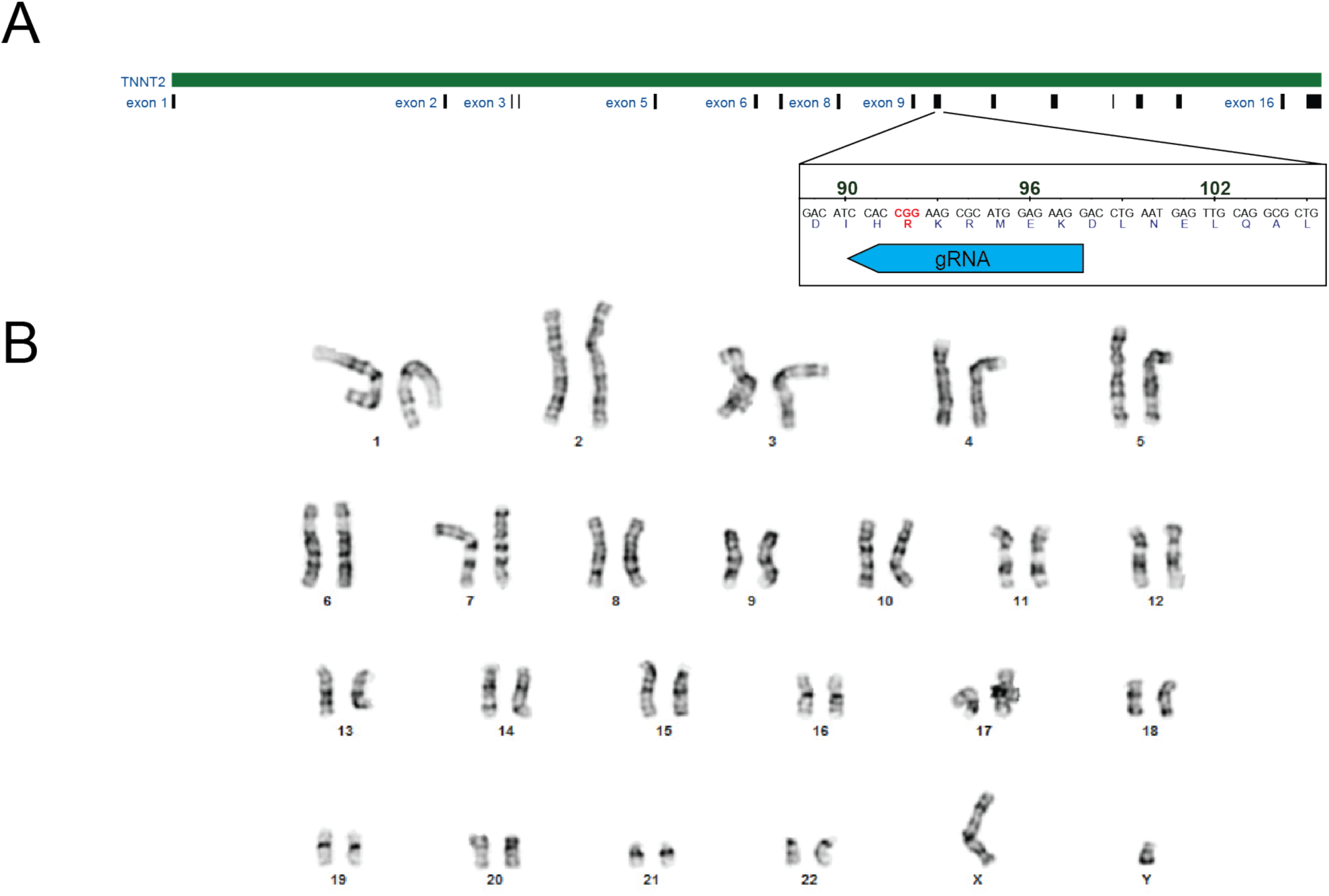
Generation of gene edited hiPSC-CMs. (A) CRISPR/Cas9 targeting of R92Q in troponin T. The R92Q mutation was added via homology directed repair. The guide RNA sequence for targeting was CCTTCTCCATGCGCTTCCGGNGG. From our screen, 21% of the cells were homozygous for the R92Q mutation (CGG -> CAA). (B) Karyotyping of R92Q gene edited cells reveals a normal karyotype.

**Supplementary Figure S2.**
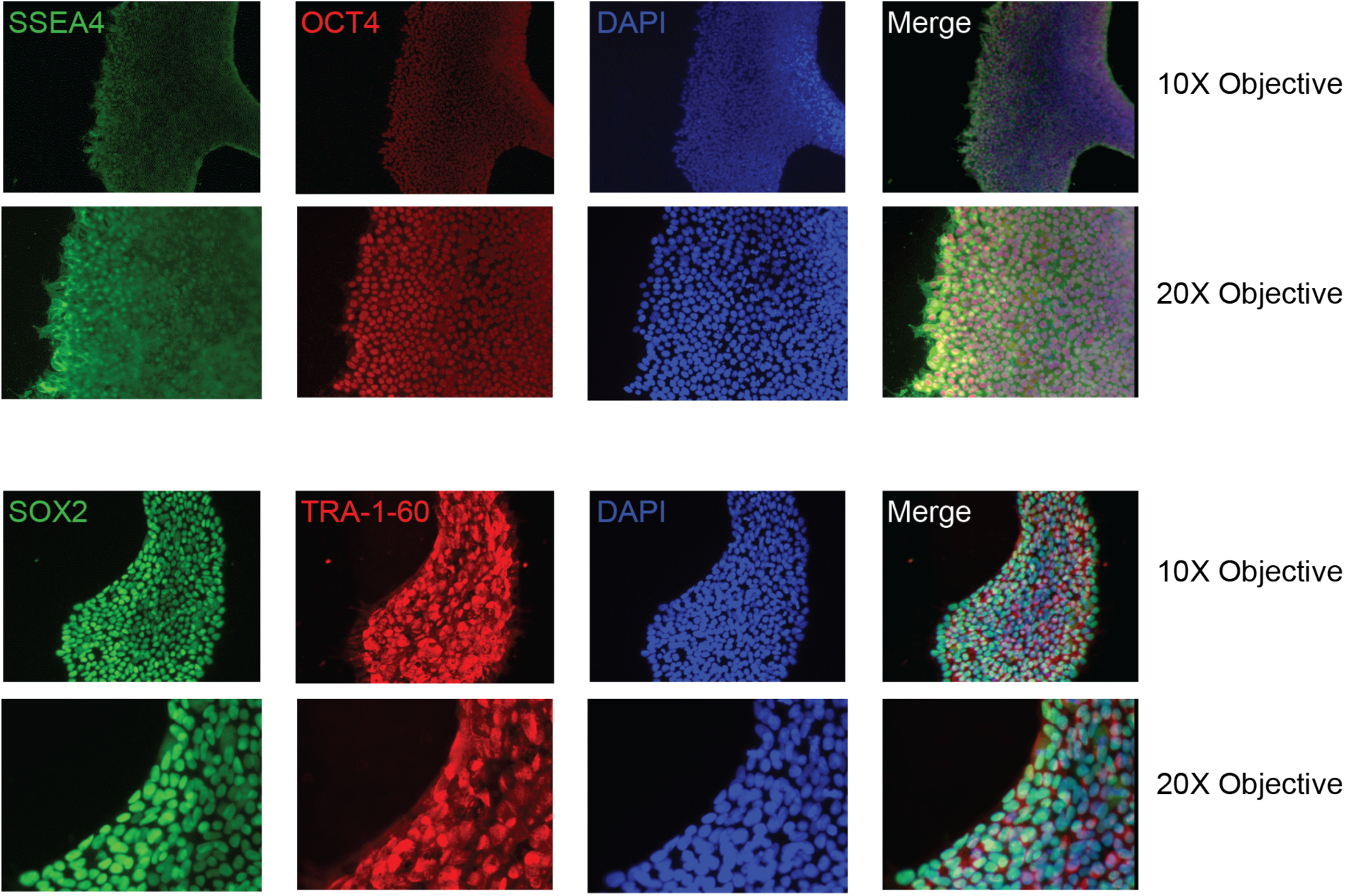
Pluripotency staining of R92Q cells. R92Q gene edited cells are pluripotent as assessed by immunofluorescence staining for the markers SSEA4, OCT4, SOX2, and TRA-1-60.

**Supplementary Figure S3.**
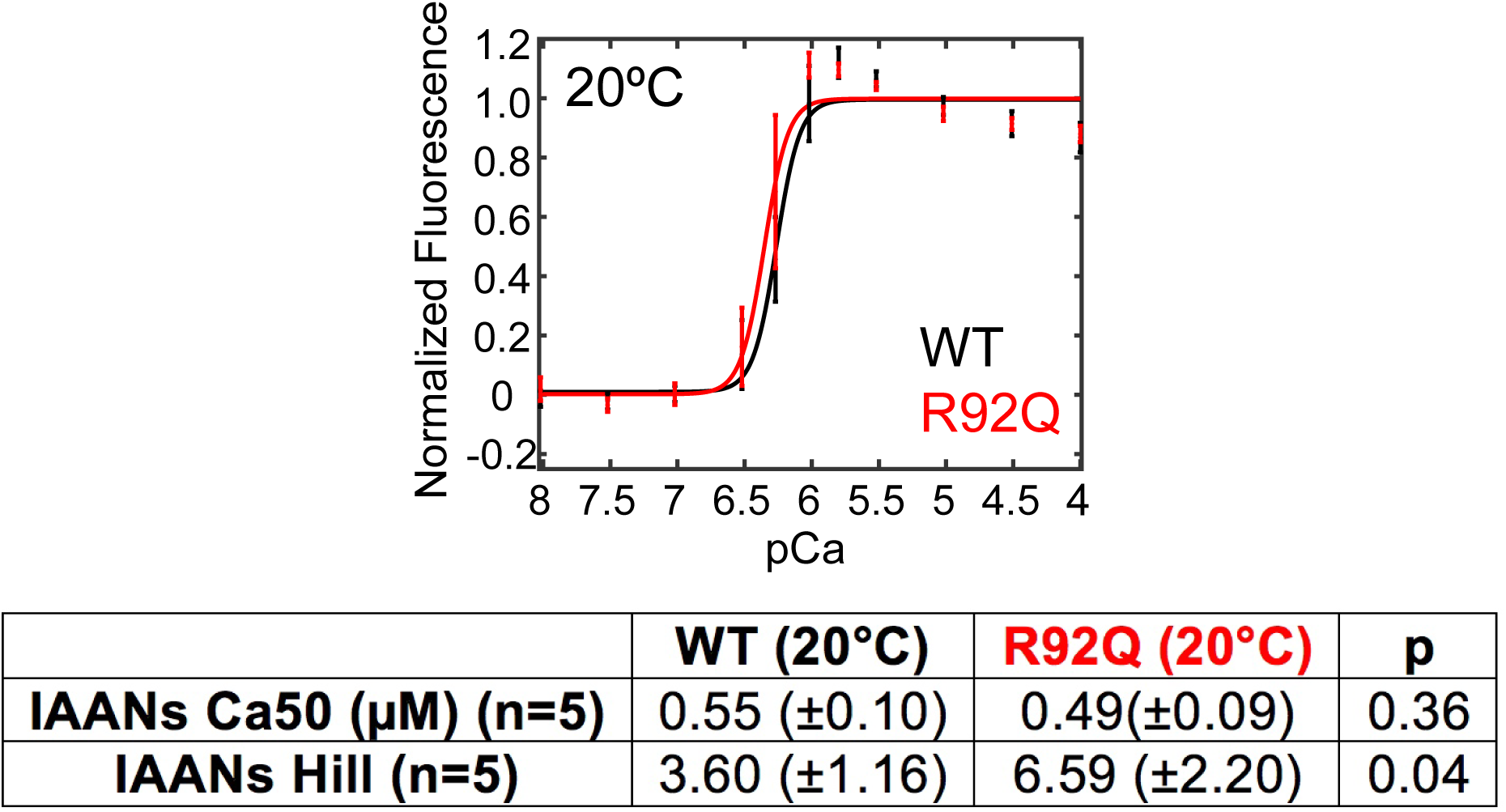
The affinity of calcium binding to the troponin complex at 20°C. IAANS-labeled troponin C was reconstituted into regulated thin filaments. Titrations with increasing calcium were conducted. Error bars show the standard deviation of 5 experiments. Values derived from fits, standard errors in the fits, and p-values are shown.

**Supplementary Table S1.** qPCR Gene Names and Primers.

| **Gene Symbol** | **Gene Name** | **Product Number** |
| --- | --- | --- |
| PLN | Phospholamban | Hs.PT.58.23189767 |
| ITPR2 | Inositol 1,4,5-trisphosphate receptor type 2 | Hs.PT.58.3479603 |
| RYR2 | Ryanodine receptor 2 | Hs.PT.58.502763 |
| CACNA1C | Calcium voltage-gated channel subunit alpha1 C | Hs.PT.58.14979004 |
| CACNA1G | Calcium voltage-gated channel subunit alpha1 G | Hs.PT.58.4441520 |
| CACNA1H | Calcium voltage-gated channel subunit alpha1 H | Hs.PT.58.814570 |
| CASQ2 | Calsequestrin 2 | Hs.PT.56a.219158 |
| ATP2A2 | ATPase sarcoplasmic/endoplasmic reticulum Ca<sup>2+</sup> transporting 2 | Hs.PT.56a.39859858.g |
| SLC8A1 | Solute carrier family 8 member A1 | Hs.PT.58.40534466 |
| CAMK2D | Calcium/calmodulin dependent protein kinase II delta | Hs.PT.56a.25723872.g |
| HPRT1 | Hypoxanthine phosphoribosyltransferase 1 | Hs.PT.58v.45621572 |
| GAPDH | Glyceraldehyde-3-phosphate dehydrogenase | Hs.PT.39a.22214836 |

**Supplementary Table S2.**
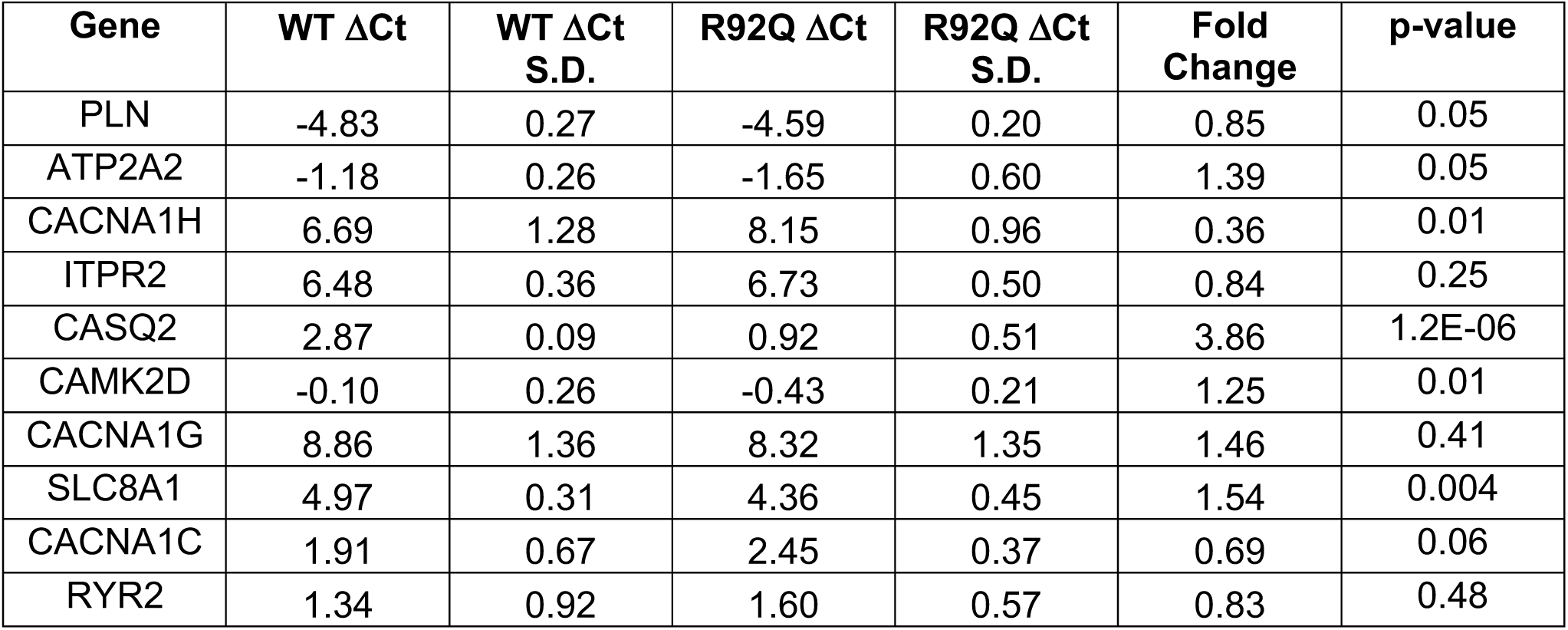
qPCR Results.

| Gene | WT ΔCt | WT ΔCt<br>S.D. | R92Q ΔCt | R92Q ΔCt<br>S.D. | Fold<br>Change | p-value |
| --- | --- | --- | --- | --- | --- | --- |
| PLN | -4.83 | 0.27 | -4.59 | 0.20 | 0.85 | 0.05 |
| ATP2A2 | -1.18 | 0.26 | -1.65 | 0.60 | 1.39 | 0.05 |
| CACNA1H | 6.69 | 1.28 | 8.15 | 0.96 | 0.36 | 0.01 |
| ITPR2 | 6.48 | 0.36 | 6.73 | 0.50 | 0.84 | 0.25 |
| CASQ2 | 2.87 | 0.09 | 0.92 | 0.51 | 3.86 | 1.2E-06 |
| CAMK2D | -0.10 | 0.26 | -0.43 | 0.21 | 1.25 | 0.01 |
| CACNA1G | 8.86 | 1.36 | 8.32 | 1.35 | 1.46 | 0.41 |
| SLC8A1 | 4.97 | 0.31 | 4.36 | 0.45 | 1.54 | 0.004 |
| CACNA1C | 1.91 | 0.67 | 2.45 | 0.37 | 0.69 | 0.06 |
| RYR2 | 1.34 | 0.92 | 1.60 | 0.57 | 0.83 | 0.48 |

**Supplementary Table S3.**
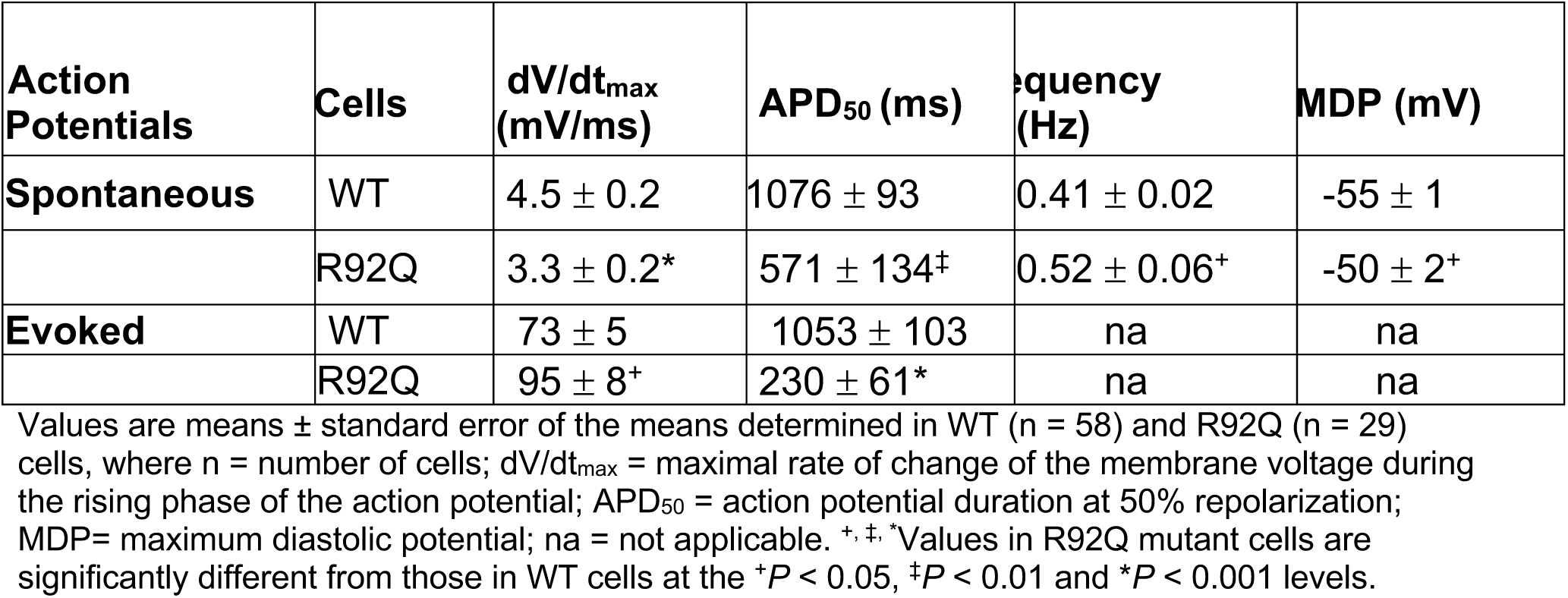
Electrophysiology Results.

| Action Potentials | Cells | dV/dt<sub>max</sub> (mV/ms) | APD<sub>50</sub> (ms) | Frequency (Hz) | MDP (mV) |
| --- | --- | --- | --- | --- | --- |
| Spontaneous | WT | 4.5 ± 0.2 | 1076 ± 93 | 0.41 ± 0.02 | -55 ± 1 |
|  | R92Q | 3.3 ± 0.2* | 571 ± 134<sup>‡</sup> | 0.52 ± 0.06<sup>+</sup> | -50 ± 2<sup>+</sup> |
| Evoked | WT | 73 ± 5 | 1053 ± 103 | na | na |
|  | R92Q | 95 ± 8<sup>+</sup> | 230 ± 61<sup>*</sup> | na | na |

Values are means ± standard error of the means determined in WT (n = 58) and R92Q (n = 29) cells, where n = number of cells; dV/dt<sub>max</sub> = maximal rate of change of the membrane voltage during the rising phase of the action potential; APD<sub>50</sub> = action potential duration at 50% repolarization; MDP= maximum diastolic potential; na = not applicable. <sup>+</sup>, <sup>‡</sup>, <sup>\*</sup>Values in R92Q mutant cells are significantly different from those in WT cells at the <sup>+</sup>P < 0.05, <sup>‡</sup>P < 0.01 and <sup>\*</sup>P < 0.001 levels.

## Notes

### Competing Interest Statement

The authors have declared no competing interest.

